# High ribulose-1,5-bisphosphate carboxylase/oxygenase content in northern diatom species

**DOI:** 10.1101/569285

**Authors:** A. C. Gerecht, G. K. Eriksen, M. Uradnikova, H. C. Eilertsen

## Abstract

Ribulose-1,5-bisphosphate carboxylase/oxygenase (RuBisCO) is a fundamental enzyme in CO_2_-fixation in photoautotrophic organisms. Nonetheless, it has been recently suggested that the contribution of this enzyme to total cellular protein is low in phytoplankton, including diatoms (< 6%). Here we show that RuBisCO content is high in some diatom species isolated from northern waters (> 69°N). Two species contained the highest RuBisCO levels ever reported for phytoplankton (36% of total protein). These high RuBisCO requirements do not increase these species’ requirements for nitrogen and do not impart a fitness disadvantage in terms of growth rate. On the contrary, high RuBisCO levels in psychrophilic diatoms may be a necessary mechanism to maintain high growth rates at low temperature at which enzymatic rates are low.

## Introduction

Ribulose-1,5-bisphosphate carboxylase/oxygenase (RuBisCO) is a fundamental enzyme in photosynthetic organisms, and has been labelled as the most abundant protein on Earth^1^. It catalyses the first step in the dark reaction of photosynthesis by binding CO_2_ to the precursor ribulose-1,5-bisphosphate. This process fixes inorganic carbon into organic matter. RuBisCO also carries out a competing oxygenase reaction, which leads to an overall loss of energy and organic matter from the cell in the process know as photorespiration. This can result in a loss of up to one third of fixed carbon in C_3_-plants^2^. The major factor determining the partitioning between carboxylase and oxygenase activity, besides the relative CO_2_ and O_2_ pressures, is the inherent ability of the enzyme to discriminate between the two substrates. This is described by the specificity factor (τ = V_C_ × K_O_ / V_O_ × K_C_) and depends on the kinetic traits of the enzyme i.e. the maximum reaction rates (V_C_, V_O_) and half-saturation constants (K_C_, K_O_) of the carboxylase and oxygenase reaction, respectively. Different lineages of photosynthetic organisms express different RuBisCO subtypes based on their evolutionary origin^3^. Data for 32 RuBisCO enzymes compiled by Badger *et al*.^4^ from cyanobacteria, algae and higher plants show a 25-fold range in the specificity factor. Whereas the kinetic traits among subtypes have been assumed to vary considerably, they are considered to be conserved within a specific subtype^5^. However, high interspecific variability in the kinetic traits of RuBisCO has recently been described for diatoms^6^.

Diatoms are important primary producers, accounting for ca. 40% of marine primary production^7^. RuBisCO in diatoms has relatively high specificity for the carboxylase reaction, similar to what is found in optimized land plants such as wheat^4, 8^. They furthermore possess an efficient carbon-concentrating mechanism (CCM) to assure high concentrations of CO_2_ near the enzyme^6^. The combination of high CO_2_-availability and an efficiently functioning RuBisCO enzyme would decrease cellular requirements for RuBisCO. In fact, diatoms have been suggested to contain low amounts of the enzyme, between 2-6% of total soluble protein^9^.

The specificity factor is furthermore influenced by environmental factors such as temperature^8^ and CO_2_-availability^10^. The specificity for the carboxylase reaction has been found to increase in four cold-water diatoms as the ambient temperature declines^8^. The oxygenase activity, on the other hand, is favoured by higher temperatures because of its higher activation energy^11^ and by the fact that the solubility of O_2_ decreases less than the solubility of CO_2_ with increasing temperature. In turn, a more efficient utilization of the CO_2_ substrate should be favoured by lower temperature, which may lower cellular requirements for RuBisCO in microalgae from cold climates. In this study, we therefore analysed the amount of RuBisCO in five marine diatoms isolated from northern waters (> 69°N), spanning a wide range of cell sizes. We also compared RuBisCO requirements to species-specific nitrogen requirements.

## Results

The growth rate (µ) was comparable among *A. longicornis*, *S. marinoi*, *C. furcellatus* and *P. glacialis* i.e. 0.25-0.28 d^−1^ (Table 1). The large diatom *C. concinnus* had doubling rates ca. one-third lower (0.18 ± 0.08 d^−1^). The cell concentrations at which the cultures were harvested (late exponential phase) depended on cell size with smaller species harvested at higher cell concentrations than larger species (Table 1). Residual nitrate concentrations were comparable for *A. longicornis*, *S. marinoi* and *P. glacialis* (Fig. 1, Table 2). However, the latter two species built up more biomass, in terms of chlorophyll *a* (chl *a*), than *A. longicornis*. *Chaetoceros furcellatus* built up a similar biomass concentration as *A. longicornis* using less nitrate, whereas the large diatom *C. concinnus* consumed ca. 7 × the amount of nitrate to build up ca. one-third of the biomass as the other species. The ratio of particulate organic carbon (POC) to nitrogen (PON; C:N-ratio) in the cells was similar among the smallest species and *P. glacialis*, whereas *C. concinnus* had a higher C:N-ratio (Table 2). *Porosira glacialis* contained high levels of RuBisCO, ca. 37% of total soluble protein (Fig. 2a, Table 2). Similarly high RuBisCO levels were measured in one of the smallest species, *S. marinoi*, whereas low levels were measured in *A. longicornis* (ca. 5%). The largest diatom *C. concinnus* had an intermediate value (ca. 9%), close to that of *C. furcellatus* (ca. 10%). When RuBisCO amounts were normalized to chl *a*, *P. glacialis* stood out as the species with highest RuBisCO levels (Fig. 2b, Table 2). These were ca. 80 × higher than those of the other species with the exception of *S. marinoi*, which had intermediate levels.

**Figure 1.**
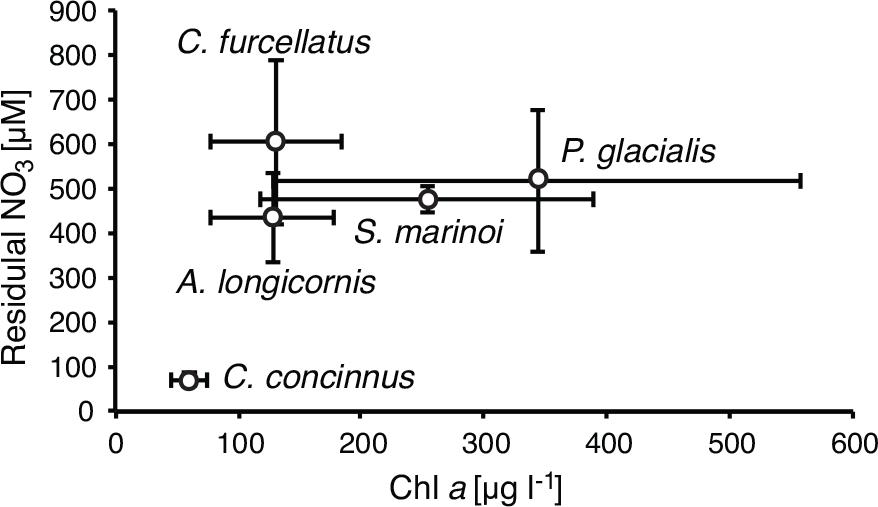
Nitrate consumption in the five diatom species. Residual nitrate concentrations plotted against final chlorophyll *a* concentrations in cultures of *A. longicornis*, *S. marinoi*, *C. furcellatus*, *P. glacialis*, and *C. concinnus*. Error bars indicate s.d. of the mean with number of replicates reported in the methods section.

**Figure 2.**
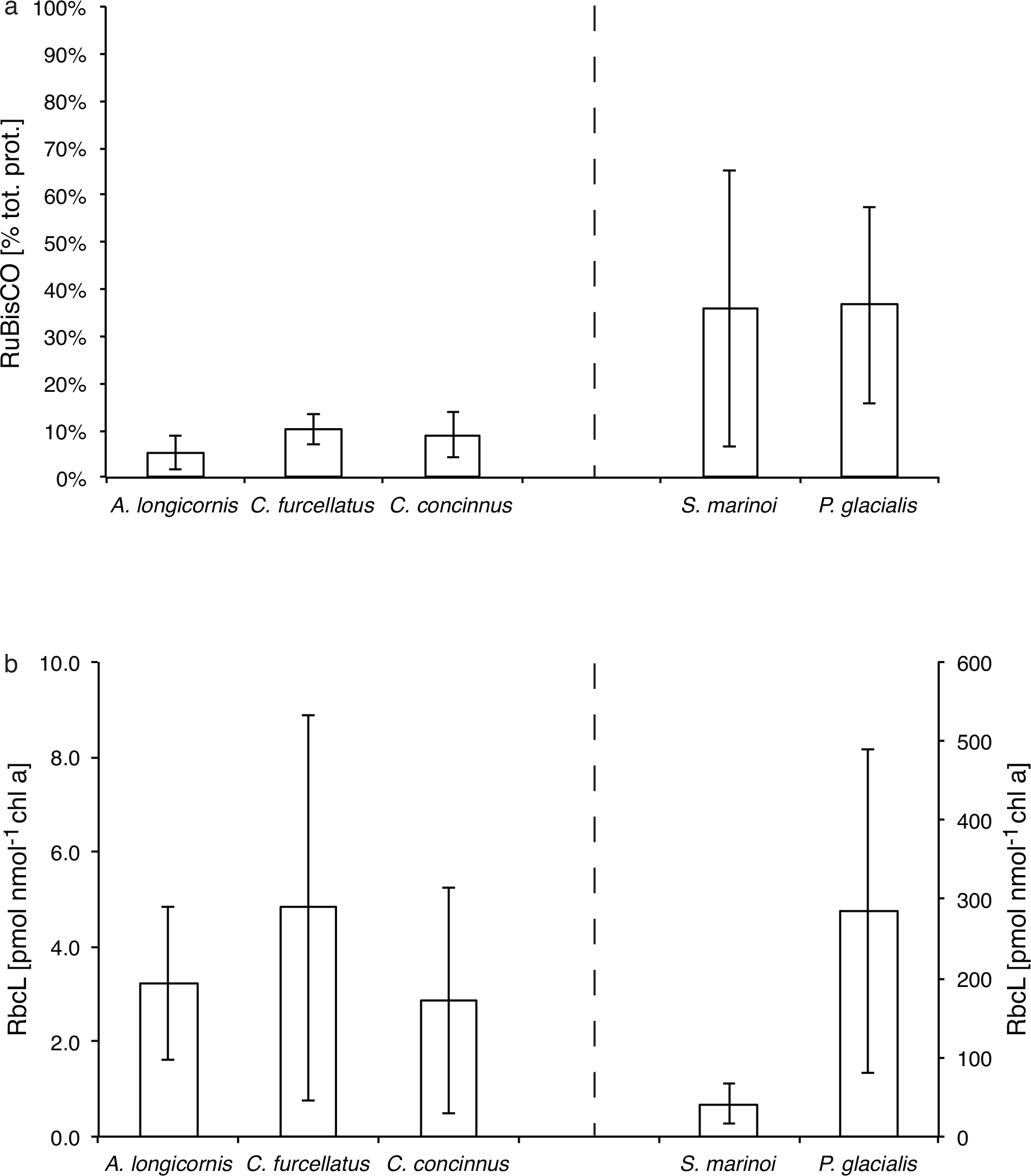
RuBisCO content in the five diatoms species. **a** RuBisCO levels in *A. longicornis*, *C. furcellatus, C. concinnus, S. marinoi*, and *P. glacialis* as percentage of total soluble protein. **b** Picomoles of RbcL normalized to chlorophyll *a*; note the different y-axes for species with low RuBisCO levels (left half) and high levels (right half). Error bars indicate s.d. of the mean with number of replicates reported in the methods section.

**Table 1.**
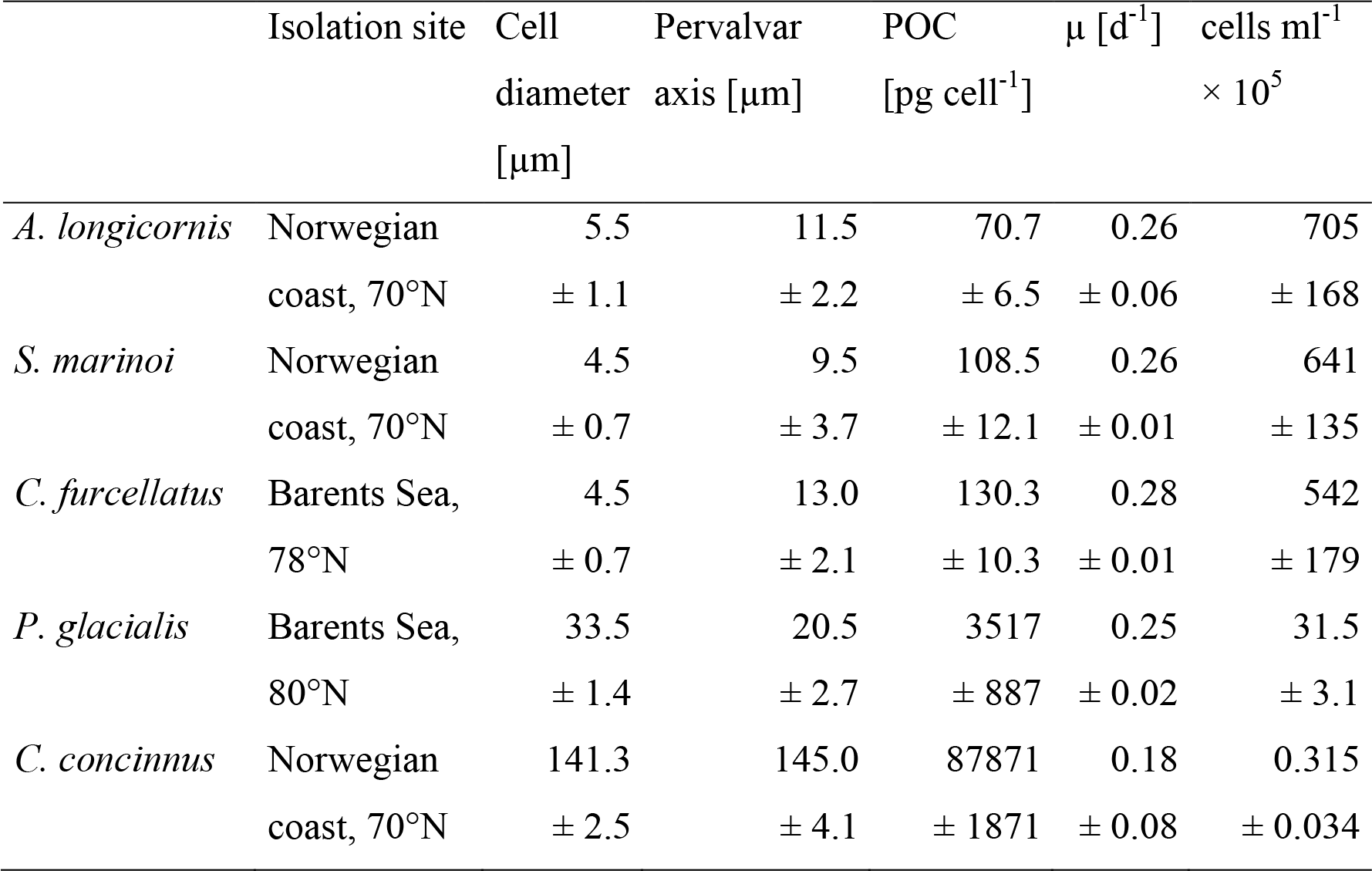
Origin, cell size, growth rate and harvest cell concentrations of the five diatom species. Means ± s.d.

**Table 2.** Nitrate consumption and RuBisCO content of the five diatom species. Means ± s.d.

|  | Residual NO<sub>3</sub><br>[μM] | Final chl *a*<br>[μg l<sup>-1</sup>] | C:N (a:a) | RuBisCO<br>[% total prot.] | RuBisCO<br>[pmol nmol<sup>-1</sup> chl *a*] |
| --- | --- | --- | --- | --- | --- |
| *A. longicornis* | 434.3<br>± 101.7 | 127.8<br>± 50.2 | 4.44<br>± 0.14 | 5.2<br>± 3.5 | 3.2<br>± 1.6 |
| *S. marinoi* | 474.5<br>± 31.0 | 253.4<br>± 136.3 | 4.17<br>± 0.22 | 35.8<br>± 29.4 | 41.2<br>± 24.7 |
| *C. furcellatus* | 604.5<br>± 185.7 | 131.0<br>± 53.3 | 4.59<br>± 0.15 | 10.3<br>± 3.3 | 4.8<br>± 4.1 |
| *P. glacialis* | 515.5<br>± 158.9 | 342.9<br>± 214.9 | 4.24<br>± 0.18 | 36.6<br>± 21.0 | 284.8<br>± 204.8 |
| *C. concinnus* | 68.8<br>± 14.6 | 60.1<br>± 14.5 | 6.75<br>± 0.75 | 9.0<br>± 4.8 | 2.8<br>± 2.4 |

## Discussion

Recently, it has been suggested that phytoplankton, including diatoms, contains low amounts of the CO_2_-fixing enzyme RuBisCO, below 6% of total soluble protein^9^. In the current study, only one diatom species, *A. longicornis*, contained similarly low amounts, whereas *C. furcellatus* and *C. concinnus* contained RuBisCO amounts within the range described previously for phytoplankton^10^. Two species, *P. glacialis* and *S. marinoi* contained strikingly high amounts of the enzyme, ca. 36% of total soluble protein. These high levels are similar to those found in land plants^1^ and have never been reported for phytoplankton. The highest previously reported levels are 23% in *Isochrysis galbana*, and most values are 5-10 times lower^10^. Considering the high interspecific variability observed in the current study, species differences may explain some of the differences among studies. Additionally, physiological differences among strains of the same species can be high in diatoms^12, 13^.

Also when RuBisCO levels were normalized to chl *a*, *S. marinoi* and *P. glacialis* stood out in terms of high RuBisCO content, whereas the other three species had values similar to those previously reported for a cyanobacteria^14^. Despite these apparently high requirements for the RuBisCO enzyme, both *S. marinoi* and *P. glacialis* showed an efficient use of nitrate. The amount of nitrate necessary to build up the same amount of biomass in these two species was actually lower than in *A. longicornis* and *C. furcellatus* and considerably lower than in the large diatom *C. concinnus*. This statement assumes that chl *a* is a cell-size and species independent proxy for biomass. However, using final POC concentrations of the culture as a biomass proxy gave qualitatively the same results i.e. lower nitrate requirements in *S. marinoi* and *P. glacialis* despite high RuBisCO content. The low C:N-ratio determined in all species except for *C. concinnus* suggests a high cellular protein content, similar to those reported for Antarctic diatoms by Young *et al*.^15^. The higher C:N-ratio in *C. concinnus* may relate to the large size of this species and higher capacity for storing C-rich compounds such as carbohydrates or lipids. There was, however, no effect of RuBisCO requirements on protein content. This may relate to a lower requirement for a CCM at high cellular RuBisCO levels and a possible trade-off between resources invested into the CCM vs. RuBisCO^5^. There are two solutions for psychrophilic organisms to overcome slow enzymatic rates at low temperatures. They can either evolve enzymes with lower thermal optima than mesophilic species or increase enzyme abundance. Descolas-Gros and de Billy^16^ determined that the activation energy for RuBisCO is the same between Antarctic and temperate diatom species, suggesting minimal cold adaptation of the enzyme. Smith and Platt^17^, on the other hand, reported differences in the temperature response between RuBisCO from Arctic and tropical phytoplankton. Similarly, Young *et al*.^15^ reported that the carboxylation rate of a mesophilic diatom decreased more strongly with decreasing temperature than that of a psychrophilic species. Both studies suggest a certain degree of cold adaptation of RuBisCO in psychrophilic diatoms. The main mechanism, however, allowing high carboxylase activity at low temperate seems to be increasing the amount of RuBisCO in the cell. The study by Young *et al*.^15^ reports RuBisCO levels of up to 23% of total protein in an Antarctic diatom bloom with similarly high levels in a psychrophilic diatom cultured at 3°C. This contrasts the study by Losh *et al*.^9^, which reports RuBisCO levels of <1% total protein in field phytoplankton samples from a temperate area. The high levels of RuBisCO reported in the current study may therefore relate to the necessity of maintaining carboxylase activity and thereby growth rate in cold-water areas and/or at low temperature in the laboratory. Similarly, psychrophilic green algae contain twice as much RuBisCO as their mesophilic counterparts^18^. Based on these studies it is tempting to draw the conclusion that our isolate of *A. longicornis* is a temperate strain transported northwards by inflowing Atlantic water, whereas the isolates of *P. glacialis* and *S. marinoi* used in this study are true cold-adapted strains. The next step is to determine to what extent RuBisCO levels are under genetic control by cultivating both meso- and psychrophilic species over a range of temperatures. Although care was taken to harvest all cultures under the same conditions, the high standard variation in *P. glacialis* and *S. marinoi*, the species with highest RuBisCO levels, may indicate a high degree of phenotypic plasticity. Preliminary data from a temperature experiment does indeed suggest lower RuBisCO levels in *S. marinoi* grow at a higher temperature (12°C; unpublished data). Similarly, Mortain-Bertrand *et al*.^19^ showed that the carboxylase activity in *S. costatum* increases from 18 to 3°C.

In conclusion, the current study provides further evidence that the strategy for psychrophilic diatoms for maintaining high growth rates at low temperature is to increase the amount of RuBisCO in the cells to allow for high carboxylase activity. It furthermore demonstrates that the requirement for RuBisCO varies strongly among diatom species. The RuBisCO levels reported here are the highest ever reported for phytoplankton. The regulation and possible constraint of these levels are areas that deserve further study.

## Methods

### Experimental setup and sample collection

The diatom species *Attheya longicornis* R. M. Crawford & C. Gardner, *Skeletonema marinoi* Sarno & Zingone, *Chaetoceros furcellatus* Yendo, *Porosira glacialis* (Grunow) Jørgensen and *Coscinodiscus concinnus* W. Smith were analysed for RuBisCO content using Western Blot. All species were either isolated from water samples or germinated from spore-containing sediment samples collected in the Barents Sea or along the coast of northern Norway (Table 1). Species were identified by a combination of morphological and molecular methods as described in Huseby^20^. Stock cultures for inoculation were kept in a climate-controlled room at 5°C and 50 µmol m^−2^ s^−1^ scalar irradiance on a 14:10 h light:dark cycle in f/10-medium^21^. Experimental cultures were grown semi-continuously as two (*C. furcellatus*, *C. concinnus*, *S. marinoi*) or three biological replicates (*A. longicornis*, *P. glacialis*) in 100 l Plexiglas cylinders under the same temperature and light conditions as stock cultures by repeatedly harvesting up to 70 l of the culture and replenishing with the same volume of nutrient-replete culture medium. This was prepared from filtered (0.22 µm), pasteurized, local seawater (Tromsø sound, 25 m depth) by adding silicate (final concentration 12.3 µM) and the commercial nutrient mixture Substral^TM^ (0.25 ml l^−1^; The Scotts Company (Nordics) A/S, Denmark). This nutrient mixture was chosen due to the need for adding economically feasible amounts of nutrients to large culture volumes^22^ and provided the following final concentrations: nitrate – 589.3 µM, ammonia – 482.1 µM, phosphate – 104.8 µM. All cultures were aerated with compressed air to avoid sedimentation and CO_2_-limitation. Growth rates (µ; Table 1) were calculated as the difference in logarithmic values of *in vitro* chl *a* between the starting and sampling day. Chl *a* was measured as three technical replicates every 2-3 days by filtering 5 mL of the culture onto a GF/C-filter, extracting the filters with 5 mL ethanol before measuring fluorescence on a TD-700 fluorometer before and after acidification. Fluorescence was converted to µg l^−1^ chl *a* according to Holm-Hansen and Riemann^23^. Cultures were harvested in late exponential phase (Table 1) by first concentrating cells onto a plankton net (mesh size 5-20 µm) before centrifuging at 3500 rpm for 5 min in a cooled centrifuge (4°C). Two technical replicates of the obtained wet pellet (ca. 400 mg each) was transferred into 2 ml Eppendorf tubes, flash-frozen in liquid N_2_ and stored at −80°C until analysis. Cell density was determined every 2-3 days and upon harvest by allowing the cells contained in 2 ml of a culture sample to settle in 2 ml Nunc culture plates before counting them under an inverted microscope. Growth rates determined from cell counts provided similar but more variable results than chl *a* data (data not shown). Cell size (diameter, pervalvar axis; Table 1) was determined on the same microscope by means of an ocular graticulate calibrated with a stage micrometer.

### Residual nitrate concentrations

Residual nitrate concentrations were measured as three technical replicates in the culture media before diluting the cultures. The cells were removed from the sample by filtering 50 ml through a GF/C filter. The samples were frozen and stored at −20°C until analysis on a Flow Solution IV analyzer, which was calibrated using reference seawater. Some of the residual nitrate concentrations were above the values calculated for initial nitrate concentrations in the culture medium based on nutrient addition (Table 2). This may be due to contamination from the natural seawater used to prepare the medium and/or the initial inoculum culture. The number of times that residual nitrate concentrations were measured in the cultures differed among species and the values plotted in Fig. 1 are based on the following number of measurements (biological replicates × dilution dates), *A. longicornis*: n=6; *C. furcellatus*: n=8; *C. concinnus*: n=4; *S. marinoi*: n=6; *P. glacialis*: n=12.

### Elemental carbon to nitrogen ratios

Three technical replicates of 50 mL each of culture was filtered onto precombusted (450°C, 8 h) GF/C filters and dried at 60°C for 24 h. The amount of POC and PON was analysed on the whole filter using a 440 elemental analyser and calibrated against standard reference material (acetanilide).

### Sample extraction and total protein analysis

To extract total soluble protein, pellet samples were suspended (1 ml: 1 g sample) in extraction buffer (50 mM MES-NaOH pH=7.0, 10 mM MgCl_2_, 1 mM EDTA, 1 mM EGTA, 1% Tween 80) containing 10 mM of the reducing agent DTT and 3% protease inhibitor cocktail. Samples were then extracted by two sonication-freeze cycles using a handheld ultrasonifier with a microtip at 20% output until the sample was just thawed before refreezing in liquid N_2_. Measurements of total soluble protein and RuBisCO levels after various sonication-freeze cycles confirmed that two cycles were sufficient to extract all soluble protein and RuBisCO from the samples, whereas additional cycles increased the likelihood of degradation (data not shown). The extract was centrifuged for 10 min at 10.000 *g* in a cooled centrifuge to remove debris and kept continuously on ice until further analysis. Total protein concentrations in the extract were measured by a reagent-compatible method (2-D Quant kit; GE Healthcare, USA).

### Western Blot analysis

0.05-2 µg of total protein were loaded onto 4-12% SDS-PAGE gels for electrophoresis using either the NuPAGE (Thermo Fisher Sci.) or Bio-Rad system (Bio-Rad, USA). For each sample two technical replicates of at least two differing protein loads were analysed. Appropriate aliquots of extract were diluted with extraction buffer before adding 1x LDS sample buffer and heating for 10 min at 70°C to denature proteins. Proteins were separated at 200 V for 30-45 min in 1x Tris/HEPES/SDS running buffer. After separation, proteins were transferred onto a PDVF membrane using either the NuPAGE wet transfer system (1 h at 60 V in Tris/glycine transfer buffer) or a semi-dry transfer system (Trans-Blot Turbo; Bio-Rad). The membrane was blocked with 5% skim milk in Tris-buffered saline/Tween 20 (TBST) buffer for 1 h at room temperature on a shaking table. The membrane was then incubated overnight at 4°C with a primary global antibody for RuBisCO (1:5000; AS03 037; Agrisera, Sweden), washed 3 times 5 min in TBST and then incubated for 1 h at room temperature with a goat anti-rabbit IgG HRP-conjugated secondary antibody (1:10000; AS09 602). The membrane was then washed as above and developed with SuperSignal West Pico Chemiluminescent substrate using an image station. A range of 3-4 standards (AS01 017S or purified diatom RuBisCO) was run alongside the samples for quantification and sample values were only considered if they fell within the standard range. As the primary antibody is designed against the large subunit of RuBisCO (RbcL), the RuBisCO amount was adjusted for the small subunit of RuBisCO (RbcS), except when using purified diatom RuBisCO as standard. RbcL amounts in the samples were quantified by direct comparison of band intensities with the standards using the system’s own software. To account for the contribution of RbcS, nanograms of RbcS were calculated using equimolar picomoles and a molecular weight of 15 kDa^9^. The number of replicates (biological × technical replicates) considered for the species was as follows, *A. longicornis*: n=13; *C. furcellatus*: n=14; *C. concinnus*: n=7; *S. marinoi*: n=9; *P. glacialis*: n=16.

## Acknowledgements

We would like to thank Svein Kristiansen for running the nitrate analyses and Paul Dubourg for carrying out the elemental CN-analyses. We would furthermore like to thank Inger Andersson for providing us with purified diatom RuBisCO.

